# hts-nim: scripting high-performance genomic analyses

**DOI:** 10.1101/261735

**Authors:** Brent S. Pedersen, Aaron R. Quinlan

## Abstract

**Motivation:** Extracting biological insight from genomic data inevitably requires custom software. In many cases, this is accomplished with scripting languages, owing to their accessibility and brevity. Unfortunately, the ease of scripting languages typically comes at a substantial performance cost that is especially acute with the scale of modern genomics datasets.

**Results:** We present *hts-nim*, a high-performance library written in the Nim programming language that provides a simple, scripting-like syntax without sacrificing performance.

**Availability:** *hts-nim* is available at https://github.com/brentp/hts-nim and the example tools are at https://github.com/brentp/hts-nim-tools both under the MIT license.

**Supplementary information:** Supplementary data are available at *Bioinformatics online*.

## 1 Introduction

For genomics applications, it can be preferable to release command-line tools over programming libraries because it limits the *surface-area* exposed to users, provides the opportunity for hidden optimizations that guarantee speed regardless of use, and eases distribution. However, command-line tools often have a limited range of functionality in an effort to minimize complexity and "feature-creep". In contrast, genomics programming libraries place a greater burden of expertise on the user, but offer flexibility in the types of analyses that be conducted. In order to provide fast, customized genomics analyses with a simple language, we expose the htslib library in the *nim* programming language in a library called *hts-nim*. *Nim* compiles to *C*, offers very good performance, and provides garbage collection, control over memory reuse, and a simple syntax that is easy for most programmers that are already familiar with common scripting languages such as Perl and Python.

We have previously published *mosdepth* (Pedersen and Quinlan, 2017) which extensively leverages *hts-nim* and is now a widely-used tool. Here we present the underlying *hts-nim* library and demonstrate how it enables the creation of fast, easy to write tools for the analysis of genomics datasets. We present an example of the syntax along with three useful command-line tools as examples. The source code for each tool is available at https://github.com/brentp/hts-nim-tools, and each script includes command-line parsing and error-handling, while also being sufficiently concise to be readable in a few minutes. The first example allows filtering a BAM/CRAM file with a simple expression language, the second counts reads in genomic regions, and the third is a quality-control tool to ensure that regions are not missing from Variant Call Format (VCF) (Danecek *et al*., 2011) files.

### 2 Approach

*hts-nim* is written in the nim programming language; low-level bindings from nim to htslib are created automatically using a tool called c2nim; then, hand-written and tested code is used for the user-exposed layer. This layer hooks into the garbage collection so that a user of *hts-nim* does not need to explicitly clean up objects or free memory as would be required in *C*. As much as possible, the interface exposed in *hts-nim* allows memory reuse to avoid pressure on the garbage collector. However, the user is also free to write code that results in more allocations for the sake of simplicity. For example, when accessing the base qualities of an alignment from a BAM file, the user passes in a *seq* variable that is filled by the *base_qualities* method in an *alignment* object; that *seq* variable is then filled inside the method and returned to the user. This allows the user to control the memory allocations, by either reusing the same container for every alignment, or allocating a new one before each call to the *base_qualities* method. The design of *hts-nim* and the *nim programming language* itself provide many opportunities for optimization like this that allow trade-offs between speed and memory.

© The Author 2018. Published by Oxford University Press. All rights reserved. For permissions, please

### 3 Examples

#### 3.1 Syntax

As an example of using the library, we present the code below that shows how a user could open, iterate, and filter a BAM.

**Listing 1.**Example syntax for BAM manipulation

~~~
import hts

var b:Bam # open a bam and index.
assert open(b, "some. bam", index=true)

for rec in b:
    if rec . qual > 10:
        echo rec . chrom , rec.start , rec.stop

# regional queries:
for rec in b. query (’ 6’ , 30867 5 , 328675 ):
  if rec.flag . proper _ pair:
    echo rec.cigar
~~~

#### 3.2 BAM/CRAM Filtering

It is common to filter alignment files using samtools (Li *et al*., 2009); here, we introduce a complementary tool built with *hts-nim*. It uses a simple expression language, *kexpr* by Heng Li, to parse and evaluate user-specified expressions. The documentation for the tool indicates the fields that are available for filtering, but briefly, flags are prefixed with *is_*, and tags are prefixed with *tag_*. This exposes a type of filtering that is not available in *samtools*. What follows is example usage searching for paired-end alignments whose ends align in the expected orientation and come from a molecule that is at least 200bp:

**Listing 2.**bam-filter example

~~~
hts -nim - tools bam - filter --threads 2 "
      is_proper_pair && insert_size > 200"
      $input_bam > $out
~~~

#### 3.3 Counting Alignments in Genomic Regions

We have previously released *mosdepth* (Pedersen and Quinlan, 2017), which leverages *hts-nim* to calculate the average per-base coverage across the genome and within a given set of regions. However, in some cases, it is preferable to know the count of reads overlapping each region rather than the average per-base coverage. The *count-reads* tool performs this operation by iterating over each read in a BAM or CRAM file while checking that the alignment meets the required flag and mapping quality constraints. The tool then tests whether the alignment overlaps the genomic regions of interest in a BED file provided on the command-line. The overlap testing is done with a *nim* library we wrote to do very fast interval lookups. Usage of this module looks like:

**Listing 3.** count-reads example

~~~
hts - nim - tool scount - reads - - threads2 - -
       mapq 10 exons.bed $input_bam > $bed
~~~

For each line in the exons.bed file, the tool will count the number of overlapping reads and report the original BED line followed by the overlap count to STDOUT. The user can also filter alignments based upon specific alignment flags, but the default excludes reads that are duplicates, failed quality-control, or secondary reads. We compared this tool to *samtools bedcov* which has a similar functionality except that it sums the total bases of coverage for each region. On an exome BAM file with 82 million reads and a BED file with about 1.2 million regions, this tool–*count-reads*–took 3 minutes and 8 seconds of CPU time while *samtools bedcov* took about 33 minutes. Simply counting the reads with *samtools view -c* takes 1 minute and 36 seconds. Rather than to compare exact times, this is to show the relative speed of *hts-nim* and to highlight the ability to create very fast, custom tools in a few lines of code.

#### 3.4 Quality Control Variant Call Files

Projects with many samples will often split the genome into regions for simple parallelization. It is possible that a few regions may result in truncated or no output because of a silent or uncaught error. This missing data can go unnoticed due the the large number of files and the complexity of processing steps. Here, we introduce a tool, *vcf-check* that takes a background VCF, e.g. from ExAC (Lek *et al*., 2016) or gnomAD (http://gnomad.broadinstitute.org/) to establish a base-line expectation of the genomic regions that are expected to have common variation. It then compares chunks of the genome from the background and the query VCF so that the user can find regions that have no representation in the query VCF but have common variation in the backgrounds. We have found this approximate metric to work well in finding regions of the genome that are lost in processing for various reasons. An example invocation of this tool looks like:

**Listing 4.**vcf-check example

~~~
vcf-check --maf 0.1 $gnomad_vcf
    $query_vcf > $missed_txt
~~~

The tab-delimited output contains the count of variants above 0.1 allele frequency for both VCFs in each region. Missing regions from the query will appear as consecutive rows with counts of 0 where the corresponding counts from the background VCF are non-zero.

## 4 Discussion

We have demonstrated the breadth of *hts-nim*’s utility by introducing a set of tools for BAM and VCF processing. These tools are available with documentation at https://github.com/brentp/ *hts-nim*-tools as a complement to the library documentation to aid users in creating their own programs. The speed and simplicity of the language combined with the utility provided by *htslib* (https://htslib.org) will make this a valuable library.

## Acknowledgements

*hts-nim* benefits from several ideas in rust-htslib which is itself an excellent genomic software library.

## Funding

This research was supported by awards to ARQ from the US National Human Genome Research Institute (NIH R01HG006693 and NIH R01HG009141), the US National Institute of General Medical Sciences (NIH R01GM124355), and the US National Cancer Institute (NIH U24CA209999).

